## Supplementary material for "Mechanics of cell integration *in vivo*": supp_information

##### I. THEORETICAL MODEL

In this work, we construct a vertex model to provide a mechanistic understanding of the process of probing and inserting by MCC cells on the overlaying epithelium. This choice is motivated by the simplicity of this class of models that involves small set of control parameters, while providing proper account of cellular shape changes together with edge and vertex dynamics. To construct the vertex model of a monolayer of cells a work functional is defined as

$$W = \frac{1}{2}\sum_{\alpha} K^{2\mathrm{D}} \left(A_{\alpha} - A_0\right)^2 + \sum_{i} \Gamma_i L_i, \tag{1}$$

where indices $\alpha$ and $i$ run over each cell and each edge, respectively. The parameter $\Gamma_i$ is the line tension within an edge $i$ with length $L_i$. Here for simplicity, we assume all cells have identical target area $A_0$ and area stiffness $K^{2\mathrm{D}}$. In case of homogeneous line tension $\Gamma$ within the tissue, the mechanical properties of the cells are controlled by a single dimensionless parameter $\gamma = \Gamma/(K^{2\mathrm{D}} A_0^{3/2})$. This definition of the work function given in Eq. 1, corresponds to a network with positive shear modulus for $\gamma > 0$, and leads to a soft network for $\gamma \leq 0$ where the ground state is degenerate under a shear deformation [1]. In this study we focus on the solid regime $\gamma > 0$. Having defined the work functional (Eq. 1), we compute the force on each vertex $\mathbf{F}_v = -\partial W/\partial \mathbf{X}_v$ and the force balance reads:

$$\frac{d\mathbf{X}_v}{dt} = \mu \mathbf{F}_v, \tag{2}$$

where $\mu$ is the mobility coefficient. To find the minimum energy configuration, not only the vertices move down the gradient according to Eq. (2), but also tissue network undergoes T1 topological transitions that are energy favorable. At each time step of the simulation we randomly pick a short edge (smaller than $\varepsilon$) and check the energy change as a result of flipping this edge $\Delta W_{T1}$. We only accept a T1 transition when $\Delta W_{T1} < 0$. Since we are interested in the force balance tissue configuration, we continue our simulation steps until the energy difference between two consecutive time steps is less than $f_{\mathrm{tol}}$.

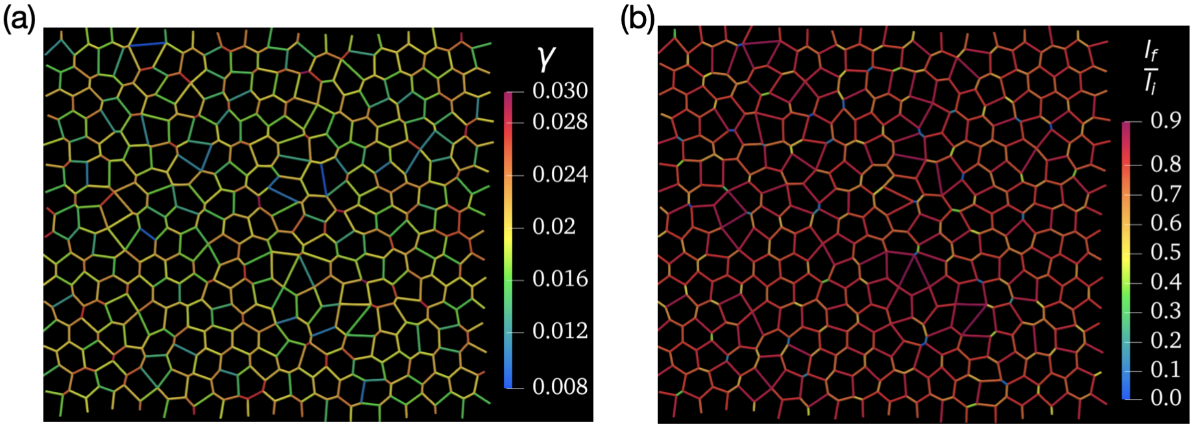


FIG. 1. **Probing the edge collapse.** a) Initial configuration before the tensional perturbation. The average line tension $<\gamma>= 0.02$. b) The line tension of each edge $i$ is perturbed $\gamma_i + 0.2 <\gamma>$. Then we find the new force balance configuration and calculate the ratio of final edge length to initial length before perturbation $l_f/l_i$.

#### II. VERTEX PROBING BY INTERCALATING CELLS

Experimental results indicate the existence of vertices with varying number of connecting edges, predominantly those with three edges (3-fold vertex) and four edges (4-fold vertex) connecting to them. To this end, we begin by simulating a monolayer of cells with heterogeneous line tension. We choose line tensions at the edges of cells within the tissue randomly from a normal distribution with the mean value of $\gamma$ and the standard deviation of $0.25 \times \gamma$. The choice of variable line tension at the cell edges ensures that both stable 3-fold and 4-fold vertices can form within the monolayer [2].

Figure 1 shows a typical configuration of cells as a result of simulating the vertex model (Eqs. 1-2). The formation of a heterogeneous cell shapes with both 3-fold and 4-fold vertices that are connected via edges of different line tension is evident. Our goal is to understand which vertices are more susceptible to be opened up by intercalating cells.

The experiments clearly demonstrate that before inserting themselves, the intercalating cells probe different vertices by extending their filopodia, making contact with the goblet cells in the monolayer, and pulling on them. In some cases this pulling continues until the intercalating cell opens up the vertex and insets itself within the monolayer, while in some other instances the intercalating cell moves to another vertex after the initial probing (see main text). As such we conjecture that the intercalating MCC cells have the ability of probing local properties of the monolayer before starting to insert themselves at particular locations. To test this, we introduce a probing mechanism to the vertex model: at each vertex an out-of-plane force of a fixed magnitude $f$ is applied, while fixing all other vertices in plane, and the out-of-plane displacement $\delta$ of the probe vertex as a result of the applied force is measured. The local measure of the stiffness at each vertex is then obtained as $\kappa_\delta = f/\delta$. Repeating this procedure for all the vertices in the monolayer gives the map of local stiffness throughout the monolayers (Fig. 3b in main text). It is clear from the stiffens map that this mechanism of out-of-plane pulling effectively probes the local tensions around different vertices distinguishing vertices with connecting edges of high line tension from those that have low tension connectivity. This is interesting, because it suggests that the pulling mechanism adapted by MCC cells could act as an effective probing of the local tension within the monolayer. Next we ask if there is any correlation between this local probing and the preferable locations of cell insertion.

#### III. MECHANISM OF VERTEX SELECTION FOR INTERCALATION

In order to explore the propensity of different vertices to intercalation, we introduce an inserting cell at each of the vertices, one at a time. The inserting cell is added with an initial area much smaller than the target area of the cell $A_{in} \ll A_0$. As such the pressure difference between the inserting cell and its neighbors works to expand the cell towards its target area. This is resisted by the cell's line tension on one hand, and is enhanced by the tension from surrounding edges at the other. As a result of this competition the inserting cell either grows to its final area or shrinks and disappears at the vertex. We distinguish between two different scenarios by defining a binary order parameter that specifies whether the cell intercalation at the vertex is successful and the MCC can integrate into the monolayer or not. Calculating the intercalation order parameter at each vertex and comparing with the probing results from the previous section we clearly observe the propensity of successful intercalation at the points where the local stiffness $\kappa_\delta$ is highest (Fig. 3b-c in main text). This is because relatively larger line tension from surrounding edges enhances the expansion of the inserting cell into the monolayer. Interestingly, we also observe that while all 4-fold vertices open up, only a fraction of 3-fold vertices with high local stiffness are chosen for intercalation. This can be further quantified by calculating the histogram of the number of 3-fold vertices that open up with respect to their local average tension. Remarkably, with increasing the line tension of the inserting cells less and less 3-fold vertices are able to open up, while still all 4-fold vertices lead to an intercalation event (Fig. 2). These results indicate that in an heterogeneous tissue a 4-fold vertex is more susceptible to be opened up by the inserting cells and as such 4-fold vertices are potential hotspots for intercalation. To understand the underlying mechanism for this distinction we next take a closer look at the instability of 3-fold and 4-fold with respect to an intercalation event.

#### IV. CRITERIA FOR VERTEX OPENING

In order to pinpoint the mechanistic basis of the easier intercalation at 4-fold vertices compared to the 3-fold vertices, here we consider a simplified setup of one intercalating cell within a pre-imposed configuration of cells with either 3-fold or 4-fold vertices. Moreover, we consider uniform constant tension across the monolayer and choose the same value of tension for the inserting cell such that the only difference between the 3-fold and 4-fold vertex is

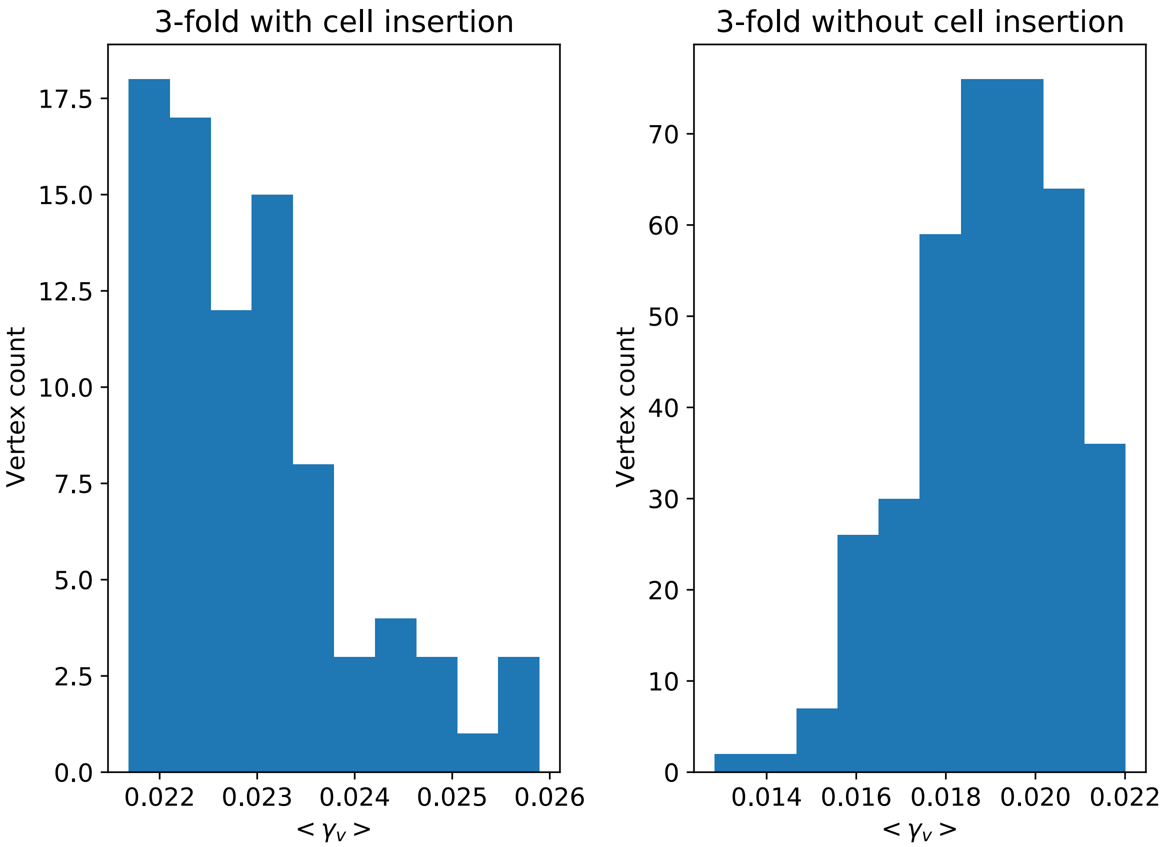


FIG. 2. Number of vertices that can open to cell with respect to their local average tension.

their configuration. We then numerically explore the possibility of intercalation for various values of tension. The results show a striking difference between the 3-fold and 4-fold vertex, demonstrating that for identical situations a 4-fold vertex opens up at a threshold value that is smaller than a 3-fold vertex (Fig. 3). To better understand this process, we next analytically investigate the criteria for vertex opening and compare our prdiction with the simplified setup discussed above.

Here we explore the intercalation of cells into the cellular network of an epithelium defined by our vertex model. We assume that the tissue can consist of three-fold, four-fold and higher fold cellular junctures. We compare the criteria for a successful cell insertion at the location of vertices with different number of connected edges.

As is depicted in Fig. 4, cells inserted at three-fold and four-fold vertices are triangular and quadrilateral respectively, and by extension $n$-gon for $n$-fold vertices. After a cell insertion, we calculate the initial forces imposed on the vertices of the inserted cells and find the criteria with which these forces are in the outward direction from the inserted cell, a necessary condition for a successful cell intercalation at the vertex.

The net force imposed on the vertex $\mathbf{r}_o$ is given by:

$$\mathbf{F}_{\mathbf{r}_o} = \frac{P_A}{2}\left[\hat{z} \times (\mathbf{u}_2 - \mathbf{u}_1)\right] + \frac{P_B}{2}\left[\hat{z} \times (\mathbf{u}_3 - \mathbf{u}_2)\right] + \frac{P_C}{2}\left[\hat{z} \times (\mathbf{u}_1 - \mathbf{u}_3)\right] \tag{3}$$

Without loss of generality we assume the vertex $\mathbf{r}_o$ at the origin of the reference frame, and $\mathbf{u}_1 - \mathbf{r}_o$ along the x-axis. Thus the opening force can be computed by projecting the net force along $\mathbf{u}_1$:

$$\mathbf{F}_{\mathbf{r}_o} \cdot \hat{u}_1 = \Gamma_1 + \Gamma_2 \hat{u}_1 \cdot \hat{u}_2 + \Gamma_3 \hat{u}_1 \cdot \hat{u}_3 + \frac{P_A}{2}\hat{z} \cdot (\mathbf{u}_2 \times \hat{u}_1) + \frac{P_B}{2}\hat{z} \cdot \left[\hat{u}_1 \times (\mathbf{u}_2 - \mathbf{u}_3)\right] + \frac{P_C}{2}\hat{z} \cdot (\hat{u}_1 \times \mathbf{u}_3) \tag{4}$$

Given that $\measuredangle \mathbf{u}_1 \mathbf{r}_o \mathbf{u}_2 = \alpha$ and $\measuredangle \mathbf{u}_1 \mathbf{r}_o \mathbf{u}_3 = \beta$, we can re-write Eq. (4) in terms of angles between the incident edges at $\mathbf{r}_o$:

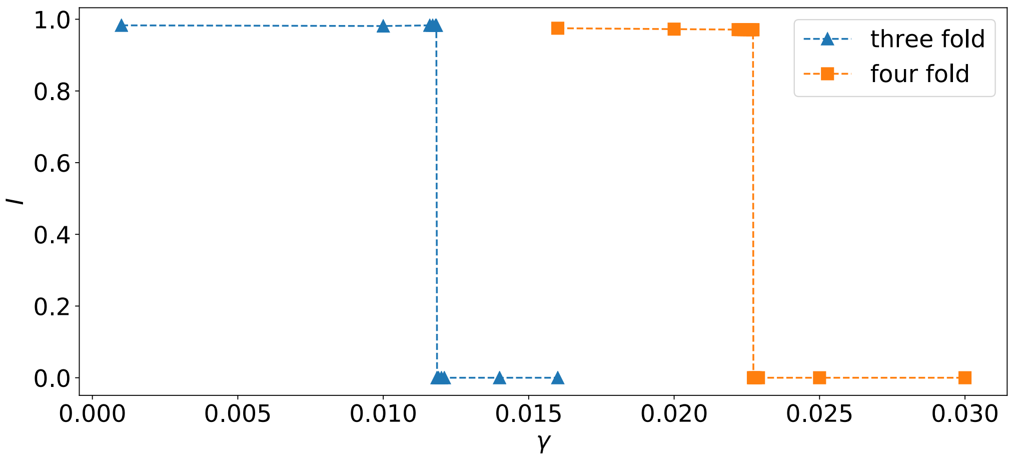


FIG. 3. Comparing the line tension criteria defined by $I$ (area of intercalated cell at force balance) for three fold and four-fold vertices, assuming a homogeneous line distribution in a honeycomb lattice.

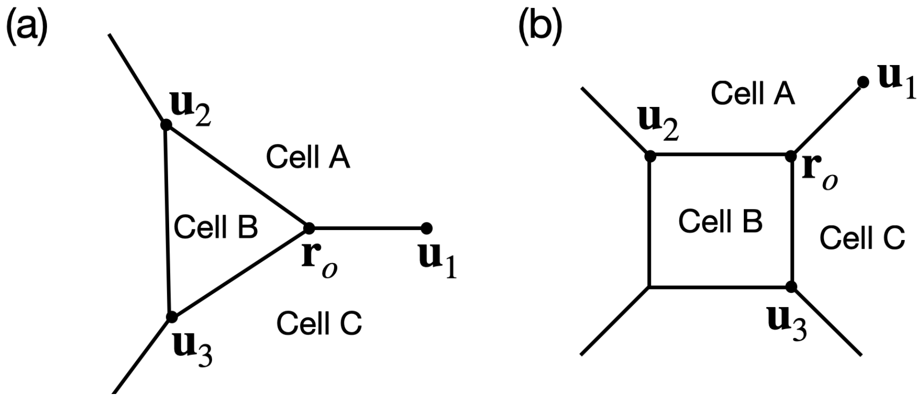


FIG. 4. a) cell insertion at three-fold vertex. b) cell insertion at four-fold vertex.

$$\mathbf{F}_{\mathbf{r}_o} \cdot \hat{u}_1 = \Gamma_1 + \Gamma_2 \cos\alpha + \Gamma_3 \cos\beta + \frac{1}{2}(P_B - P_A)|\mathbf{u}_2| \sin\alpha + \frac{1}{2}(P_C - P_B)|\mathbf{u}_3| \sin\beta \tag{5}$$

The necessary condition for a successful cell insertion reads:

$$\mathbf{F}_{\mathbf{r}_o} \cdot \hat{u}_1 > 0. \tag{6}$$

To simplify this criteria, we consider completely symmetric configuration where the $n$-fold vertex opens into a regular $n$-gon (i.e. the three-fold vertex opens into a equilateral triangles and the four-fold vertex opens to a square). Further assuming that cells $A$ and $C$ have the same pressure equal to $P_N$, and defining the pressure of cell $i$ as $P_i = -K^{2\mathrm{D}}(A_i - A_0)$, the opening force on an n-fold vertex simplifies to:

$$\mathbf{F}_{\mathbf{r}_o} . \hat{u}_1 = \Gamma_1 - (\Gamma_2 + \Gamma_3) \sin(\frac{\pi}{n}) + \frac{K^{2\mathrm{D}}}{2}(A_N - A_B) \cos(\frac{\pi}{n})(|\mathbf{u}_2| + |\mathbf{u}_3|) \tag{7}$$

We can simplify this even further, by assuming $|\mathbf{u}_2| = |\mathbf{u}_3| = \epsilon\sqrt{A_0}$, and equal line tension for edges of the inserted cell $\Gamma_2 = \Gamma_3 = \Gamma_n$. The opening force normalized by line tension $f_o = \mathbf{F}_{\mathbf{r}_o} \cdot \hat{u}_1/\Gamma_1$:

$$f_o = 1 - 2\frac{\gamma_n}{\gamma_1} \sin(\frac{\pi}{n}) + \frac{a_N - a_B}{\gamma_1} \epsilon \cos(\frac{\pi}{n}), \tag{8}$$

where the dimensionless quantities are $a_i = A_i/A_0$, and $\gamma_i = \Gamma_i/(K^{2\mathrm{D}}A_0^{3/2})$. For positive values of line tension parameter $\Gamma_1$, the opening condition given in Eq. 6 reads $f_o > 0$.

Figure (5) shows the cell intercalation regimes based on Eq 8 only for three-fold and fold-vertices.

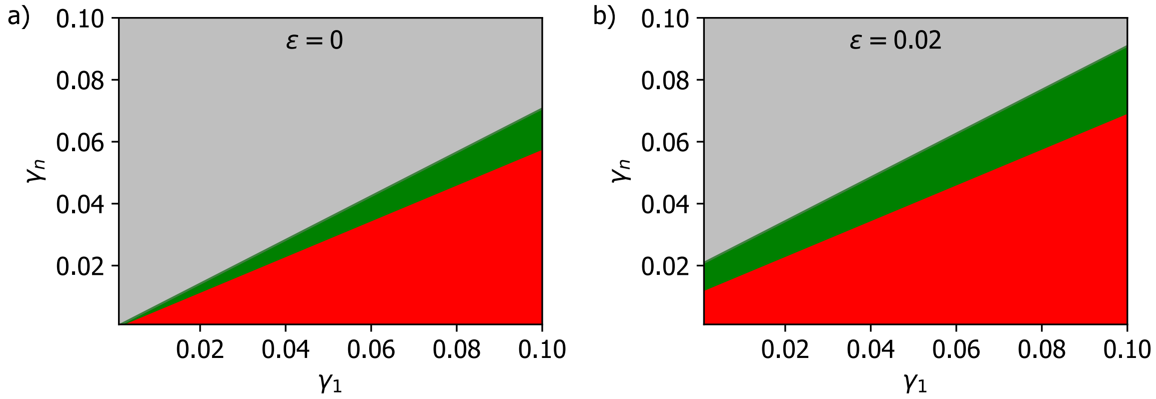


FIG. 5. Stability diagram for the intercalation at 3-fold and 4-fold vertices. Gray: neither three-fold nor four-fold allows a cell intercalation. Green: Only four-fold vertices allow cell intercalation. Red: Both three-fold and four-fold vertices allow for cell intercalation. a) Infinitesimal size of inserting cells. b) Finite size of inserting cells.

#### V. PROBING AND COLLAPSING EDGES FOR INTERCALATION

Not only the intercalating cells prefer inserting themselves at 4-fold vertices rather than 3-fold ones, but also we explore the possibility of them actively rearranging the epithelial layer to form 4-fold vertices. This is possible for example by probing and pulling at two vertices of the same edge and closing that edge such that two adjacent 3-fold vertices collapse into a 4-fold vertex, which we showed in previous section is a preferential cite for the cell to insert itself. To investigate this interesting mechanism, we conduct numerical experiment in which we probe the response of edges - one at a time - to external contraction. This is informed from the experimental observations that show intercalating MCC cells can pull simultaneously on the two vertices of the same edge. We thus conjecture that depending on the length and the tension of the edge, this simultaneous pulling can induce an edge collapse actively remodeling the epithelial layer to form preferential 4-fold vertices. The results of these simulations are summarized in Fig. 1. In order to quantify the edge fate under tensional perturbation we define $l_f/l_i$ as the order parameter characterising the ratio of the particular edge's length after applying the perturbation to that edge's initial length. As such, if an edge collapses under tensional perturbation and a 4-fold vertex is formed, the order parameter is zero, $l_f/l_i = 0$, and it is finite otherwise. The simulation results show that the edge collapse and the subsequent 4-fold vertex formation can only occur for edges with sufficiently short length and sufficiently large tension. This can be more clearly seen from the stability-diagram of $l_f/l_i$ in the length-tension phase space clearly demonstrating the propensity of edge collapse for small initial length and high tensions within the edge (Fig. 4n). Taken together, these results show that when intercalating MCCs pull on two vertices of an edge in the goblet layer, they can indeed induce a re-arrangement in the goblet layer by closing that edge and forming a 4-fold vertex, which in turn is a preferential site for the MCCs to insert themselves. Therefore, not only MCCs take advantage of existing 4-fold vertices in the goblet layer, they can even create their own 4-fold vertex from an edge, if the edge is sufficiently short and its tension is sufficiently large.

[1] R. Farhadifar, J.-C. Röper, B. Aigouy, S. Eaton, and F. Jülicher, Current Biology **17**, 2095 (2007).
[2] M. A. Spencer, Z. Jabeen, and D. K. Lubensky, The European Physical Journal E **40**, 2 (2017).

### SUPPLEMENTARY FIGURES

#### FIGURE S1

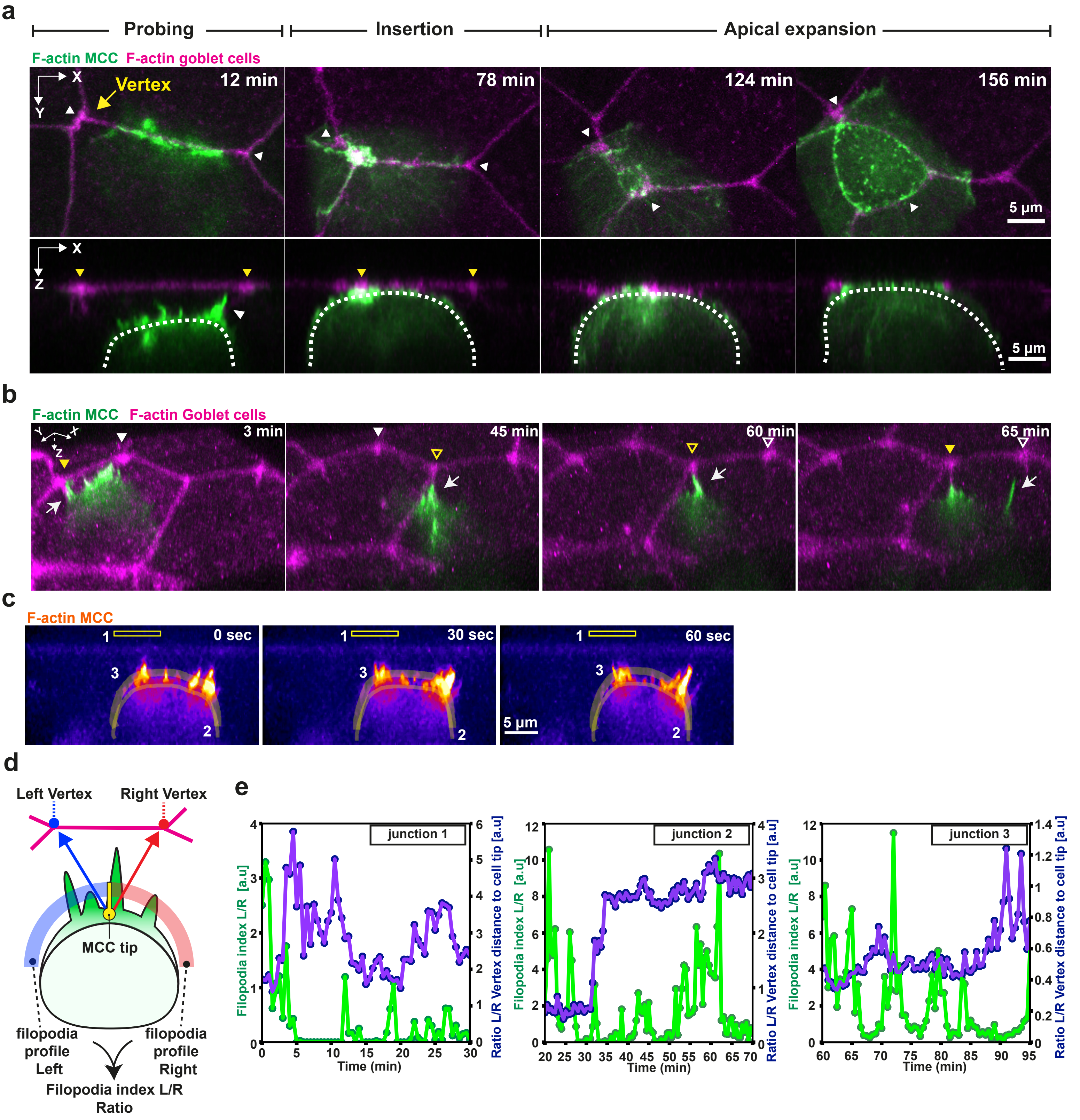

**Supplementary figure 1: Probing is the first stage of MCC intercalation.** Dynamics of MCC probing. Intercalating MCCs are visualized by expression of MCC specific α-tubulin:LifeAct-GFP (green) while overlying goblet cells are visualized by expression of goblet cell specific nectin promoter driven Utrophin-RFP (magenta). **a,** Snapshots of the different steps of MCC radial intercalation. White arrowheads mark direction for reslicing. In the corresponding orthogonal (XZ) projections yellow arrowheads mark vertices and white arrow marks F-actin filopodium. White dotted lines outline the cell contour. Scale bars: 5 µm. **b,** 3D rendering of intercalating MCC interacting with different vertices (marked by arrowheads)(MCC from Fig. 1b-d). **e,** Representation of filopodia dynamics quantification. Different regions of interest (ROIs) corresponding to the background (1), cortex (2) and filopodia (3) are drawn and used to extract F-actin intensity. Scale bar: 5 µm. **d,** Schematics describing protrusion analysis (continued). Protruding MCCs are divided into two segments (left in blue, right in red) and the filopodia dynamics is calculated for each half. **f,** Filopodia dynamics during lateral movement. Distance ratio between left/right vertex (purple) and filopodia ratio between left/right vertex (green).

### FIGURE S2

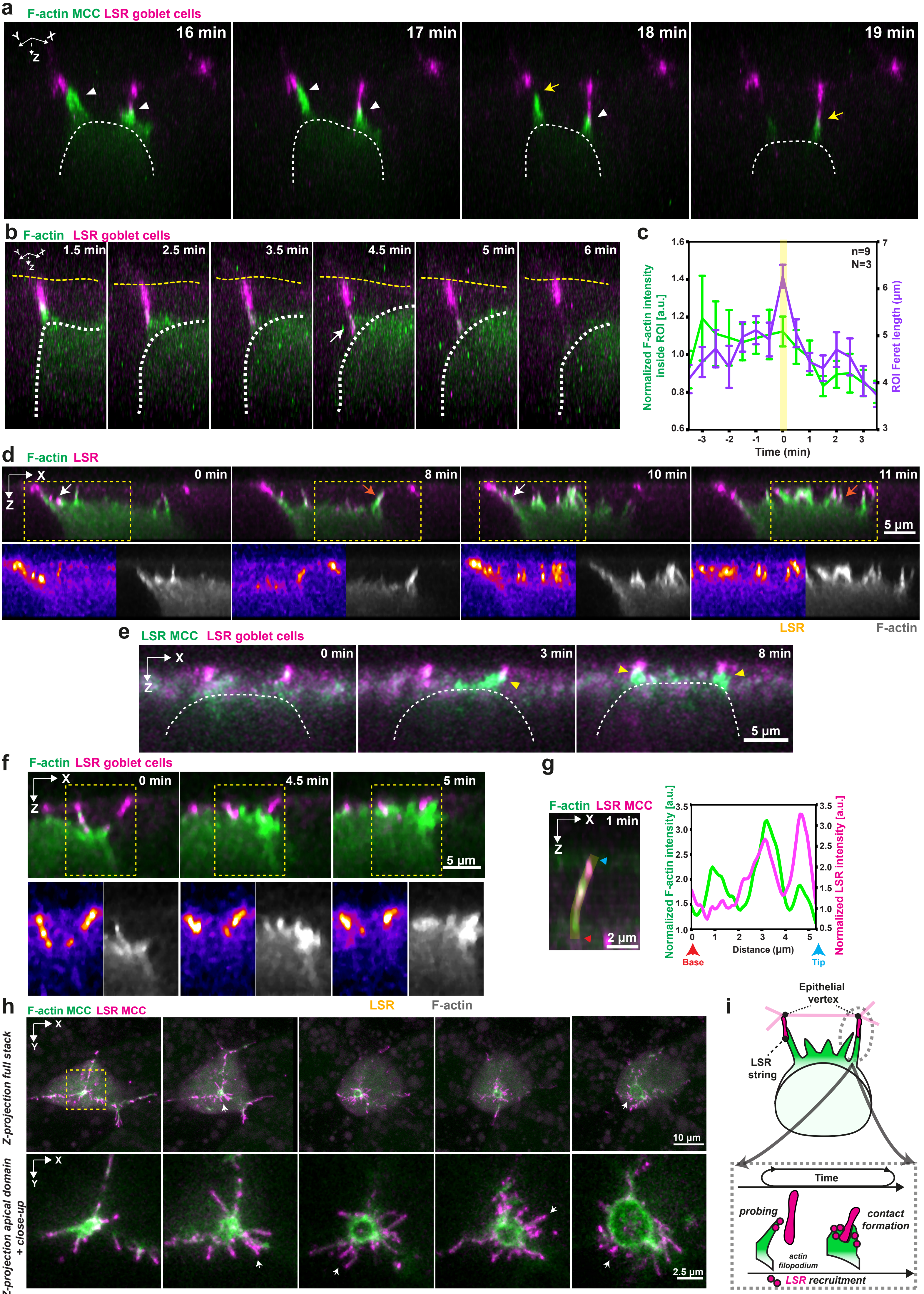
a
F-actin MCC LSR goblet cells
16 min
17 min
18 min
19 min
b
F-actin LSR goblet cells
1.5 min
2.5 min
3.5 min
4.5 min
5 min
6 min
c
n=9
N=3
Normalized F-actin intensity inside ROI [a.u.]
ROI Feret length (µm)
Time (min)
d
F-actin LSR
0 min
8 min
10 min
11 min
5 µm
LSR
F-actin
e
LSR MCC LSR goblet cells
0 min
3 min
8 min
5 µm
f
F-actin LSR goblet cells
0 min
4.5 min
5 min
5 µm
g
F-actin LSR MCC
1 min
2 µm
Normalized F-actin intensity [a.u.]
Normalized LSR intensity [a.u.]
Distance (µm)
Base
Tip
LSR
F-actin
h
F-actin MCC LSR MCC
Z-projection full stack
10 µm
Z-projection apical domain + close-up
2.5 µm
i
Epithelial vertex
LSR string
Time
probing
actin filopodium
contact formation
LSR recruitment

**Supplementary figure 2: LSR is required for interaction with the vertices. a,** 3D rendering of intercalating MCC during the probing phase. The intercalating cell (F-actin in green) uses filopodia to form close attachments to the vertices of epithelial neighbors (LSR-GFP in magenta). White arrowheads depict filopodia-goblet cell contact and yellow arrows depict contact retraction. **b**, 3D rendering of filopodia pulling on vertex (marked by white arrow, from Fig. 2e). White dotted line outlines the MCC contour and yellow dotted line outlines the top of the superficial epithelium. **c**, Epithelial vertex pulling quantification. Average MCC F-actin intensity (green) and vertex length (purple) during pulling and retraction (pulled). T=0 marks the vertex length maxima. Error bars represent SEM. **d,** Orthogonal (XZ) projections of intercalating MCC (F-actin in green) expressing LSR-3xGFP (magenta). LSR is recruited to the contact points between the MCC and the vertex (white arrow, t=0 min and t= 10 min) and to filopodia (orange arrow, t=8 min and t=11 min)). Yellow boxes mark insets with separate channels. Scale bar: 5 µm **e,** Orthogonal (XZ) projections of intercalating MCC expressing LSR-GFP (green) interacting with vertices labeled with LSR-RFP (magenta). Yellow arrowheads mark the putative homophilic LSR-LSR contacts. Scale bar: 5 µm **f,** Orthogonal (XZ) projections of intercalating MCC expressing Lifeact-RFP (green) interacting with vertices labeled with LSR-GFP (magenta). Insets depict F-actin recruitment in the MCC to stabilize the new contact. Scale bar: 5 µm **g,** Plot profile of F-actin and LSR along filopodium (from Fig. 2d). Normalized intensity values are plotted from base to tip. Scale bar: 2.5 µm. **h,** Image sequence of LSR-overexpressing MCC. Ectopic protrusions are labeled with white arrows. Top row: Full projection. Yellow box marks inset for bottom row, close-up on the apical domain. **i**, Schematics depicting LSR recruitment to the leading edge of intercalating MCC.

### FIGURE S3

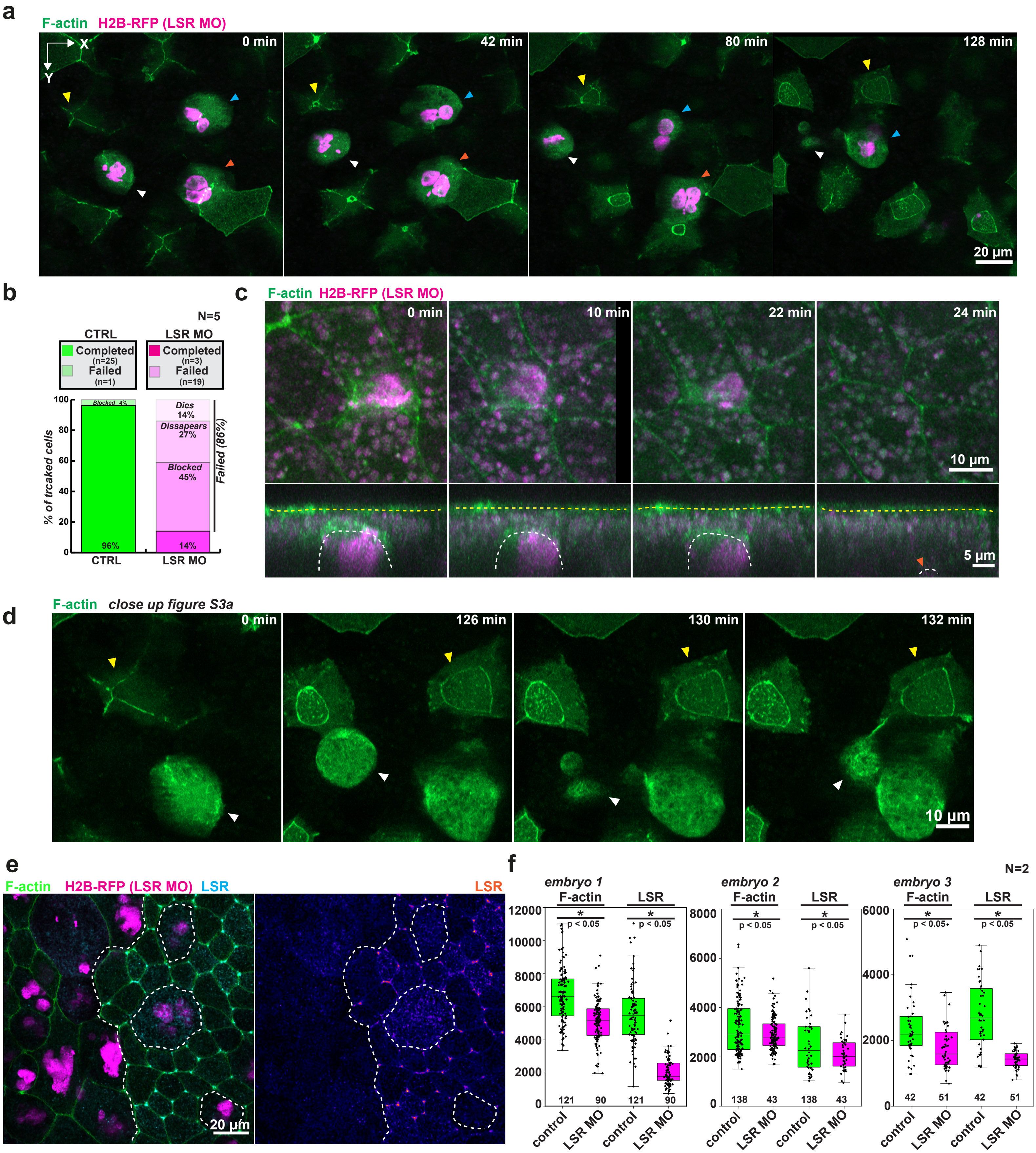
a
F-actin H2B-RFP (LSR MO)
X
Y
0 min
42 min
80 min
128 min
20 µm
b
N=5
CTRL
LSR MO
Completed (n=25)
Failed (n=1)
Completed (n=3)
Failed (n=19)
% of trcaked cells
100
80
60
40
20
0
Blocked 4%
96%
Dies 14%
Dissapears 27%
Blocked 45%
14%
Failed (86%)
CTRL
LSR MO
c
F-actin H2B-RFP (LSR MO)
0 min
10 min
22 min
24 min
10 µm
5 µm
d
F-actin close up figure S3a
0 min
126 min
130 min
132 min
10 µm
e
F-actin H2B-RFP (LSR MO) LSR
LSR
20 µm
f
N=2
embryo 1
embryo 2
embryo 3
F-actin
LSR
*
p < 0.05
12000
10000
8000
6000
4000
2000
0
121
90
121
90
8000
6000
4000
2000
0
138
43
138
43
6000
4000
2000
0
42
51
42
51
control
LSR MO

**Supplementary figure 3: LSR controls MCC intercalation. a,e,** Control and LSR depleted cells (marked with H2B-RFP, magenta) express LifeAct-GFP (green). Image sequence of control MCC (yellow arrowhead) and LSR depleted MCCs. LSR depleted cells fail to intercalate (blocked, cyan arrowhead), migrate back inside the tissue (disappear, orange arrowhead) or undergo cell death (dies, white arrowhead) during cell intercalation. Scale bar: 20 µm. **b,** Quantification of intercalation success rates. LSR depleted cells are either blocked, disappear, or dying (nWT= 26 cells, nLSRMO = 22 cells, N=5 experiments). **c,** Image sequence depicting disappearance of LSR depleted MCC (marked by orange arrowhead). **d,** Image sequence depicting cell death of LSR depleted MCC (marked by white arrowhead). A control MCC is marked by a yellow arrowhead. Scale bar: 10 µm. **e,** Immunofluorescence image of LSR stained embryo. White dotted line marks the boundaries between control and LSR depleted goblet cells (marked by H2B-RFP, in magenta). **f**, Distribution of F-actin and LSR intensities in control (marked in green, total n=301 vertices, 3 embryos, N=2 experiments) and LSR depleted cells (marked in magenta, total n=183 vertices, 3 embryos, N=2 experiments).

FIGURE S4

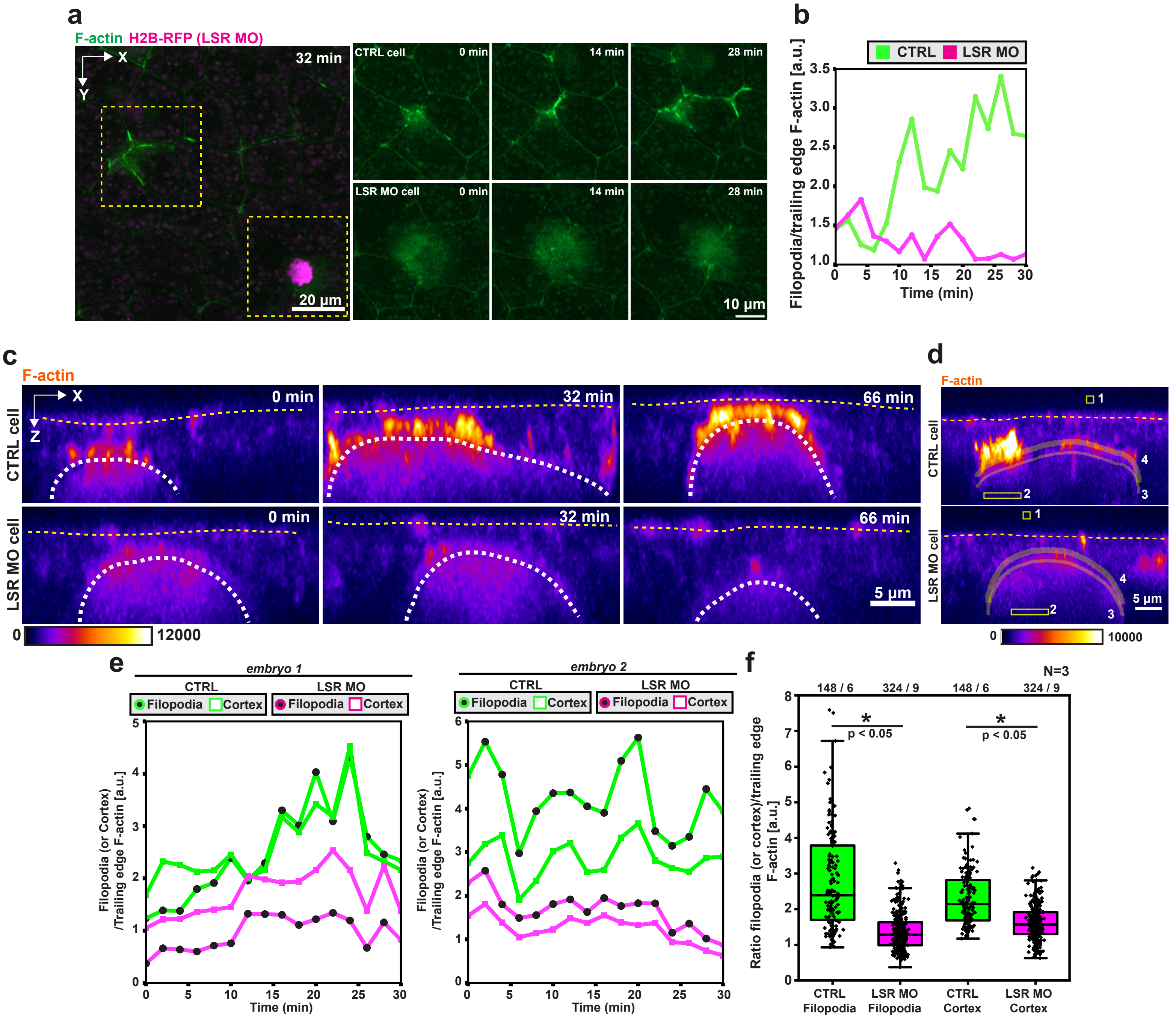
a
F-actin H2B-RFP (LSR MO)
32 min
X
Y
20 µm
CTRL cell
0 min
14 min
28 min
LSR MO cell
0 min
14 min
28 min
10 µm
b
CTRL LSR MO
Filopodia/trailing edge F-actin [a.u.]
Time (min)
c
F-actin
CTRL cell
LSR MO cell
0 min
32 min
66 min
5 µm
0
12000
d
F-actin
CTRL cell
LSR MO cell
5 µm
0
10000
e
embryo 1
embryo 2
CTRL
LSR MO
Filopodia
Cortex
Filopodia (or Cortex) /Trailing edge F-actin [a.u.]
Time (min)
f
N=3
148 / 6
324 / 9
148 / 6
324 / 9
p < 0.05
p < 0.05
Ratio filopodia (or cortex)/trailing edge F-actin [a.u.]
CTRL Filopodia
LSR MO Filopodia
CTRL Cortex
LSR MO Cortex

**Supplementary figure 4: LSR controls F-actin dynamics in MCCs.**

**a,** F-Actin (green) dynamics in control and LSR depleted marked with H2B-RFP (magenta). Scale bar: 20 µm. Yellow boxes mark insets for control and LSR depleted MCC. Scale bar for insets: 10 µm. **b,** Quantification of F-actin accumulation at the leading edge of control (green) and LSR MO (magenta) cells from g. **c,** Orthogonal (XZ) projections used for quantitative analysis of F-actin dynamics in control and LSR depleted cells expressing LifeAct-GFP (pseudo-colored in fire). Mosaic cells were extracted from the same embryo and image grey values were adjusted to the maximum and minima for comparison. **d,** Representation of the ROIs extracted for quantitative analysis: 1 - background, 2 - trailing edge, 3 - cortex and 4 - filopodia. **e,** Normalized F-actin intensity at the cortex (squares) and filopodia (circles) in control (green) and LSR depleted cells (magenta) **f,** Box plots representing distribution of F-actin intensities at the cortex and filopodia of control (n=148 timepoints pooled from 6 cells, N=3 experiments) and LSR depleted cells (n=324 timepoints pooled from 9 cells, N=3 experiments).

### FIGURE S5

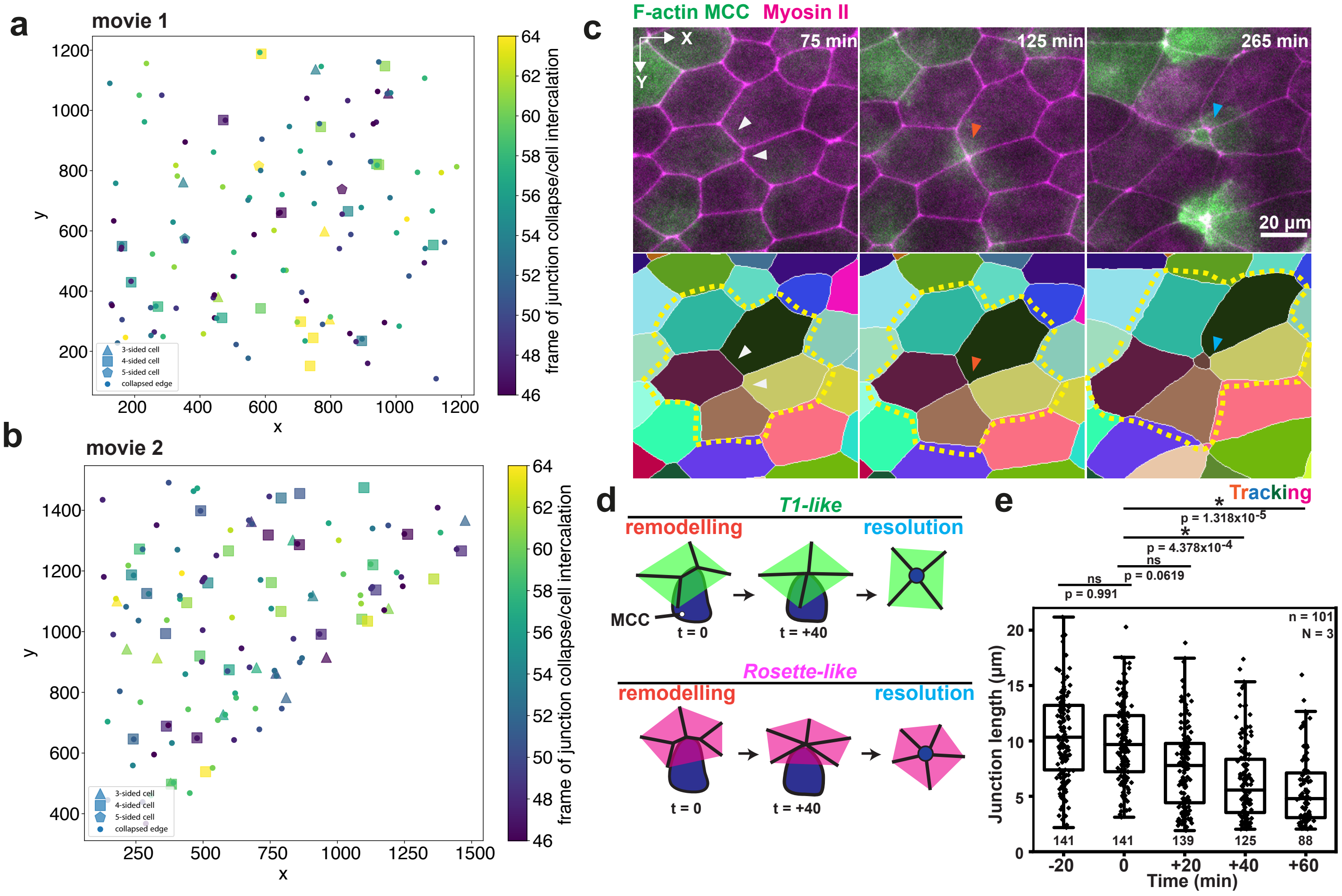
a
movie 1
3-sided cell
4-sided cell
5-sided cell
collapsed edge
x
y
frame of junction collapse/cell intercalation
b
movie 2
c
F-actin MCC Myosin II
X
Y
75 min
125 min
265 min
20 μm
d
T1-like
remodelling
resolution
MCC
t = 0
t = +40
Rosette-like
remodelling
resolution
e
Tracking
p = 1.318x10-5
p = 4.378x10-4
ns
p = 0.0619
ns
p = 0.991
n = 101
N = 3
Junction length (μm)
Time (min)
141
141
139
125
88

**Supplementary figure 5: MCC intercalation happens simultaneously with junction remodelling. a-b,** Spatial correlation between edge collapse (circles) and MCC intercalation (polygons), colored by the time of occurrence. **c,** Higher-fold MCC intercalation. White arrowhead mark initial junction layout, orange arrowhead mark junction collapse and cyan arrowhead marks apical expansion. Yellow outline depicts rosette formation. Scale bar: 20 μm **d,** Schematics representing intercalation at higher fold vertices, which is divided in a remodelling phase and a resolution phase. Intercalation at a 4-fold vertex resembles a T1 event (T1-like) whereas 5-fold intercalation or higher resembles the formation of a rosette (Rosette-like). **e,** Junction length distribution involved in forming higher-order vertices. T=0 marks the onset of MCC intercalation (junction number > 88, n=101 cells from N=3 embryos).

### FIGURE S6

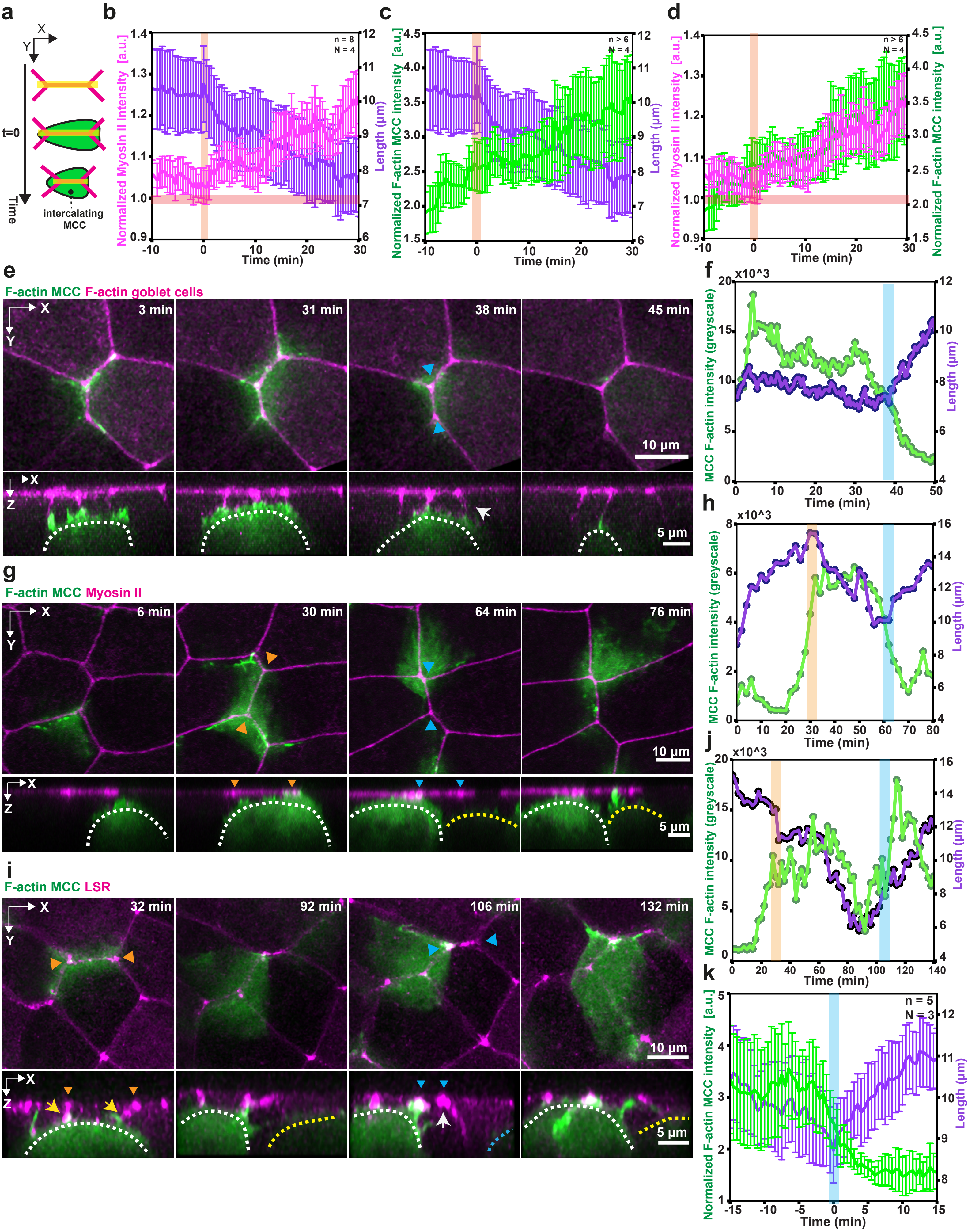
a
b
c
d
e
F-actin MCC F-actin goblet cells
3 min
31 min
38 min
45 min
10 µm
5 µm
f
g
F-actin MCC Myosin II
6 min
30 min
64 min
76 min
10 µm
5 µm
h
i
F-actin MCC LSR
32 min
92 min
106 min
132 min
10 µm
5 µm
j
k
t=0
Time
intercalating MCC
Normalized Myosin II intensity [a.u.]
Length (µm)
Normalized F-actin MCC intensity [a.u.]
MCC F-actin intensity (greyscale)
Time (min)
n = 8
N = 4
n > 6
N = 4
n = 5
N = 3
x10^3

**Supplementary figure 6: MCCs pull on neighboring vertices to induce remodelling.** Orange bars and arrowheads mark onset of junction collapse, cyan bars and arrowheads mark onset of junction retraction. **a,** Schematics representing junction (in magenta) remodelling quantification during MCC intercalation (in green). **b,** Average normalized junctional myosin-II intensity (magenta) and length (purple) before and after collapse (marked by t=0)(n=8 junctions from N=4 experiments). **c,** Average normalized MCC F-actin intensity (green) and length (purple) before and after collapse (marked by t=0)(n>6 junctions from N=4 experiments). **d,** Normalized junctional myosin-II intensity (magenta) and normalized MCC F-actin intensity (green) during collapse (marked by t=0)(n>6 junctions from N=4 experiments). **e,g,i,** Image sequence depicting junction retraction after MCC (in green) losses of contact (marked by cyan arrowheads) with the vertex. White dotted line marks cell of interest, yellow dotted line marks competing MCC. Yellow arrow marks contact with the vertex and white arrow marks loss of contact with the vertex. **f,h,j,** Corresponding MCC F-actin intensity in grey values (green) and length (purple) during junction retraction. **k,** Average normalized MCC F-actin intensity (green) and length (purple) before and after retraction (marked by t=0)(n=5 junctions from N=3 experiments). Error bars represent SEM.

### FIGURE S7

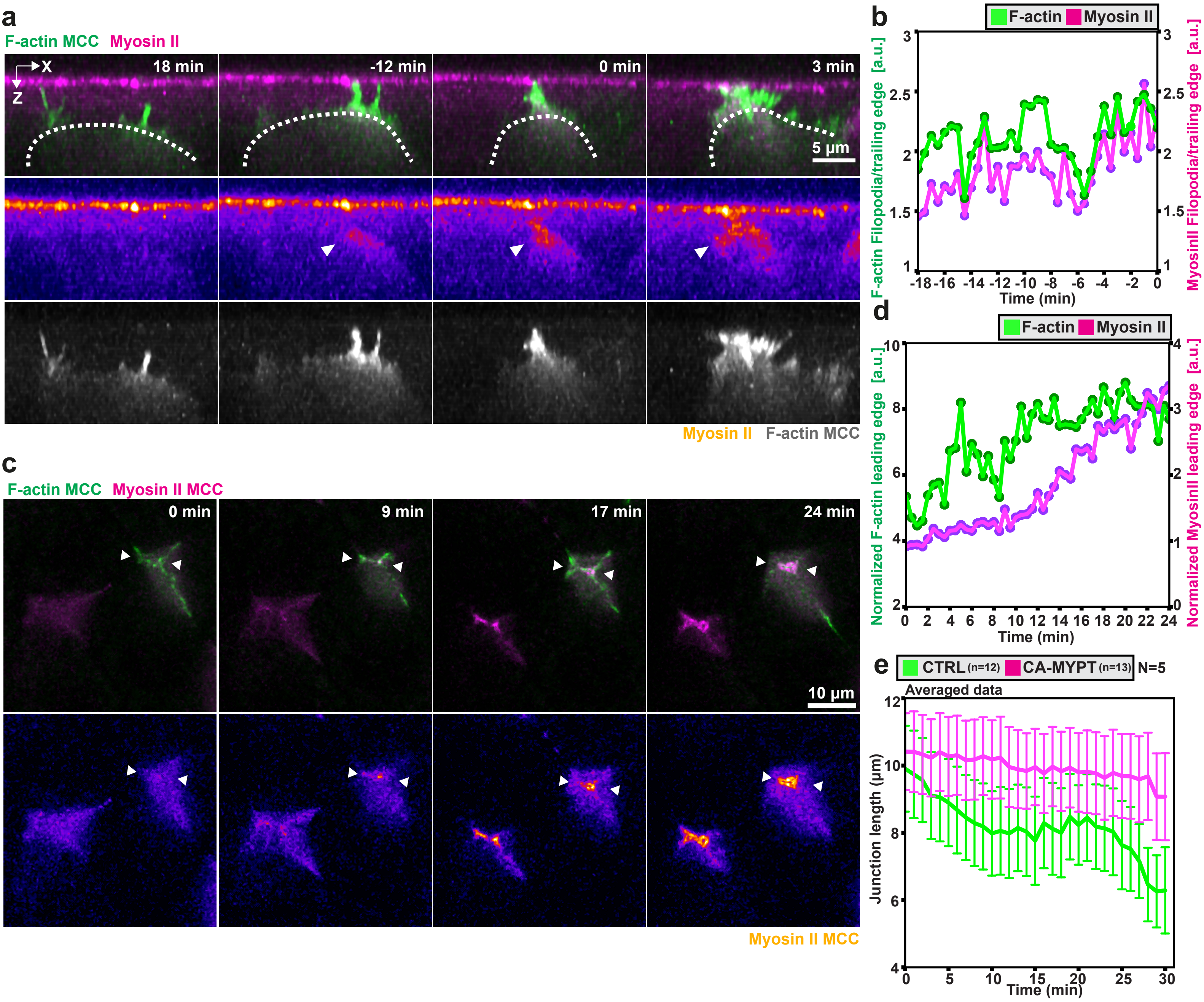
a
F-actin MCC Myosin II
X
Z
18 min
-12 min
0 min
3 min
5 µm
Myosin II F-actin MCC
b
F-actin Myosin II
F-actin Filopodia/trailing edge [a.u.]
MyosinII Filopodia/trailing edge [a.u.]
Time (min)
c
F-actin MCC Myosin II MCC
0 min
9 min
17 min
24 min
10 µm
Myosin II MCC
d
F-actin Myosin II
Normalized F-actin leading edge [a.u.]
Normalized MyosinII leading edge [a.u.]
Time (min)
e
CTRL (n=12) CA-MYPT (n=13) N=5
Averaged data
Junction length (µm)
Time (min)

**Supplementary figure 7: Intercalating MCCs use Myosin II during intercalation and remodelling.** **a,** Orthogonal (XZ) projections of myosin recruitment in MCC during intercalation. Intercalating MCC and Myosin-II are labeled with α-tubulin LifeAct-GFP (green) and the myosin nanobody SF9-3xGFP (magenta), respectively. Scale bar: 5 µm. t=0 marks the last frame before the cell contacts the epithelial vertex. **b,** Normalized myosin-II intensity (magenta) and normalized MCC F-actin intensity (green) at the leading edge of intercalating MCC. **c,** Image sequence depicting myosin recruitment in MCC during junction remodelling. White arrowheads depict myosin accumulation **d,** Normalized myosin-II intensity (magenta) and normalized MCC F-actin intensity (green) at the leading edge of inserted MCC. **e,** Average junction length for control and CAMYPT-overexpressing MCC (nWT= 12 junctions, nCAMYPT = 13 junctions, N=5 experiments).

### SUPPLEMENTARY VIDEOS

**Video Legends**

***Supplementary Video 1:*** 3D rendering of intercalating MCC interacting with epithelial vertices. MCC expressing α-tubulin:LifeAct-GFP (green) and goblet cells expressing nectin:utrophin-RFP (magenta).

***Supplementary Video 2:*** 3D plots for filopodia dynamics. Relative position of F-actin protrusions (magenta) extended by an intercalating MCC (cyan) and the overlaying epithelial vertices (horizontal tracks, color-coded for distance) during lateral movement.

***Supplementary Video 3:*** 3D rendering of MCC interacting with epithelial vertices. Intercalating MCC expressing α-tubulin:LifeAct-RFP (green) and goblet cells expressing nectin:LSR-GFP (magenta).

***Supplementary Video 4:*** LSR localizes to filopodia tips. Intercalating MCC expressing α-tubulin:LifeAct-RFP (green) and α-tubulin:LSR-GFP (magenta). Scale bar: 2 µm

***Supplementary Video 5:*** LSR depletion blocks MCC intercalation. Intercalating control and LSR MO MCCs expressing LifeAct-GFP (green). LSR depleted cells are marked with H2B-RFP (magenta). Scale bar: 20 µm

***Supplementary Video 6:*** LSR overexpression induces the formation of ectopic filopodia. Intercalating MCC expressing α-tubulin:LifeAct-RFP (green) and α-tubulin:LSR-GFP (magenta). Scale bar: 5 µm

***Supplementary Video 7:*** 3D rendering of filopodia pulling on epithelial vertices. Intercalating MCC expressing α-tubulin:LifeAct-RFP (green) and goblet cells expressing LSR-3xGFP (magenta).

***Supplementary Video 8:*** Orthogonal view of filopodia pulling on epithelial vertices. Intercalating MCC expressing α-tubulin:LifeAct-RFP (green) and goblet cells expressing nectin:LSR-GFP (magenta). Scale bar: 5 µm

***Supplementary Video 9:*** Rosette-like structure formation during MCC intercalation. Intercalating MCCs expressing α-tubulin:LifeAct-RFP (green) and goblet cells expressing SF9-3xGFP (magenta). Scale bar: 20 μm

***Supplementary Video 10:*** Junction remodelling during MCC intercalation. Intercalating MCC expressing α-tubulin:LifeAct-RFP (green) and goblet cells expressing SF9-3xGFP (magenta). Scale bar: 5 μm

***Supplementary Video 11:*** Junction retraction after contact loss. Intercalating MCC expressing α-tubulin:LifeAct-RFP (green) and goblet cells expressing SF9-3xGFP (magenta). Scale bar: 10 μm

***Supplementary Video 12:*** Orthogonal view of MCC vertex retraction after contact loss. Intercalating MCC expressing α-tubulin:LifeAct-RFP (green) and goblet cells expressing LSR-3xGFP (magenta). Scale bar: 5 μm

***Supplementary Video 13:*** Myosin II is recruited to the leading edge of intercalating MCCs. Goblet cells and MCC expressing SF9-3xGFP (magenta). Intercalating MCC expresses MCC marker α-tubulin:LifeAct-RFP (green). Scale bar: 10 μm

***Supplementary Video 14:*** Myosin II downregulation blocks junction remodelling and cell intercalation. Goblet cells and CA-MYPT overexpressing MCC expressing LF-GFP (green). CA-MYPT overexpressing MCC is labeled with H2B-RFP (magenta). Scale bar: 10 μm
